# Cobalt(III) Schiff Base complexes stabilize non-fibrillar amyloid-β aggregates with reduced toxicity

**DOI:** 10.1101/2020.05.12.091405

**Authors:** K. F. Roberts, C. R. Brue, A. Preston, D. Baxter, E. Herzog, E. Varelas, T. J. Meade

## Abstract

The aggregation of Aβ is believed to be foundational to the pathogenesis of Alzheimer’s disease (AD). *In vitro* aggregation kinetics have been shown to correlate with rates of disease progression in both AD patients and animal models, thus proving to be a useful metric for testing Aβ-targeted therapeutics. Here we present evidence of Cobalt(III) Schiff base complex (Co(III)-sb) modulation of Aβ aggregation kinetics by a variety of complementary techniques. These include Thioflavin T (ThT) fluorescence, circular dichroism (CD) spectroscopy, transmission electron microscopy (TEM), and atomic force microscopy (AFM). Our data was fitted to kinetic rate laws using a mathematical model developed by Knowles et al. in order to extract mechanistic information about the effect of Co(III)-sb on aggregation kinetics. Our analysis revealed that Co(III)-sb significantly decreases the kinetic parameter k_+,_ and significantly increases the polymerization rate k_n_, suggesting that Co(III)-sb causes Aβ to rapidly form stable oligomeric species that are unable to elongate into mature fibrils. This result was corroborated by TEM and AFM of Aβ aggregates *in vitro*. We also demonstrate that Aβ aggregate mixtures produced in the presence of Co(III)-sb exhibit decreased cytotoxicity compared to untreated samples.

**Statement of Significance:** Amyloid-β is thought to be a key mediator in the pathology of Alzheimer’s disease, yet its precise mechanisms of toxicity are poorly understood. The interaction of Aβ with endogenous metal ions via its N terminal Histidine residues has been shown to alter the peptide’s aggregation and toxicity. As such, metal-based complexes have been developed both as therapeutic agents as well as tools for investigating the role of metal binding in the pathogenesis of AD. This work expands on our previous studies developing Cobalt(III) Schiff base complexes as amyloid inhibitors. Here we demonstrate effective inhibition of aggregation by various complementary modalities. Additionally we show that Co(III)-sb reduces the toxicity of Aβ aggregates to cells in culture.

## Introduction

The amyloid-β (Aβ) peptide is a key mediator in the etiology of Alzheimer’s disease (AD), the most common form of age-related dementia and sixth leading cause of death in the US.(1, 2) Pathologically, AD is characterized by both the formation of amyloid plaques composed of Aβ and intracellular aggregates composed of the hyperphosphorylated microtubule-associated protein, tau. Disruptions in tau processing are common to many neurodegenerative disorders and, in the case of AD, generally believed to be downstream of Aβ toxicity.(2-6) The precise mechanism by which these pathological changes induce synaptic dysfunction and neurodegeneration remains unknown.

Aβ is produced extracellularly following the sequential cleavage of the transmembrane protein APP by β- and γ-secretase. The two primary isoforms *in vivo* are Aβ_40_ and Aβ_42_, which are present in an approximately 9:1 ratio in cerebrospinal fluid.(2, 7) Initially monomeric and unstructured, Aβ pathologically undergoes amyloid aggregation in AD. It is unclear what triggers the aggregation of the soluble innocuous metabolite Aβ.

Amyloid aggregation follows a nucleation-dependent polymerization reaction wherein monomers associate into small oligomers. The oligomers then act as nuclei for further polymerization. The formation of nuclei is slow and termed the “lag phase” of aggregation. Once a critical nucleus is formed, the proteins rapidly aggregate into larger-order aggregates and eventually fibrils rapidly during the polymerization phase. Equilibrium is eventually reached between the various aggregate species. In addition to this primary nucleation process, various secondary nucleation mechanisms have been identified and mathematically described.(8, 9) Significant work by Knowles et al. has resulted in a system of rate laws which model kinetic aggregation data and relate the curves to underlying microscopic processes.(10-12)

While the utility of *in vitro* amyloid aggregation has been broadly demonstrated, assays measuring protein aggregation are subject to a host of confounding variables such as peptide concentration, pH, ionic strength, temperature, agitation conditions, and molecular crowding.(13, 14) With so many potential confounding variables, it is to be expected that there is staggering variability in published aggregation data.(15) Although studies measuring *in vitro* amyloid aggregation have significant limitations, it is generally accepted that in order to overcome the limitations of individual assays, it is best to confirm results using a variety of complementary assays.

Extensive research has identified the three N-terminal histidine residues of Aβ (positions 6, 13, 14) as modulators of the peptide’s aggregation propensity and toxicity.(16-26) These residues form a metal ion binding site which has been shown to interact with endogenous copper and zinc ions.(16, 20, 24) Coordination to these residues alters the peptides geometry which leads to changes in the aggregation properties. Additionally, coordination to copper increases Aβ’s neurotoxicity, while coordination to zinc decreases toxicity, likely secondary to a reactive oxygen species mediated mechanism. Therefore, development of coordination complexes that target these N-terminal His residues has been an active area of research.

**Scheme 1:**
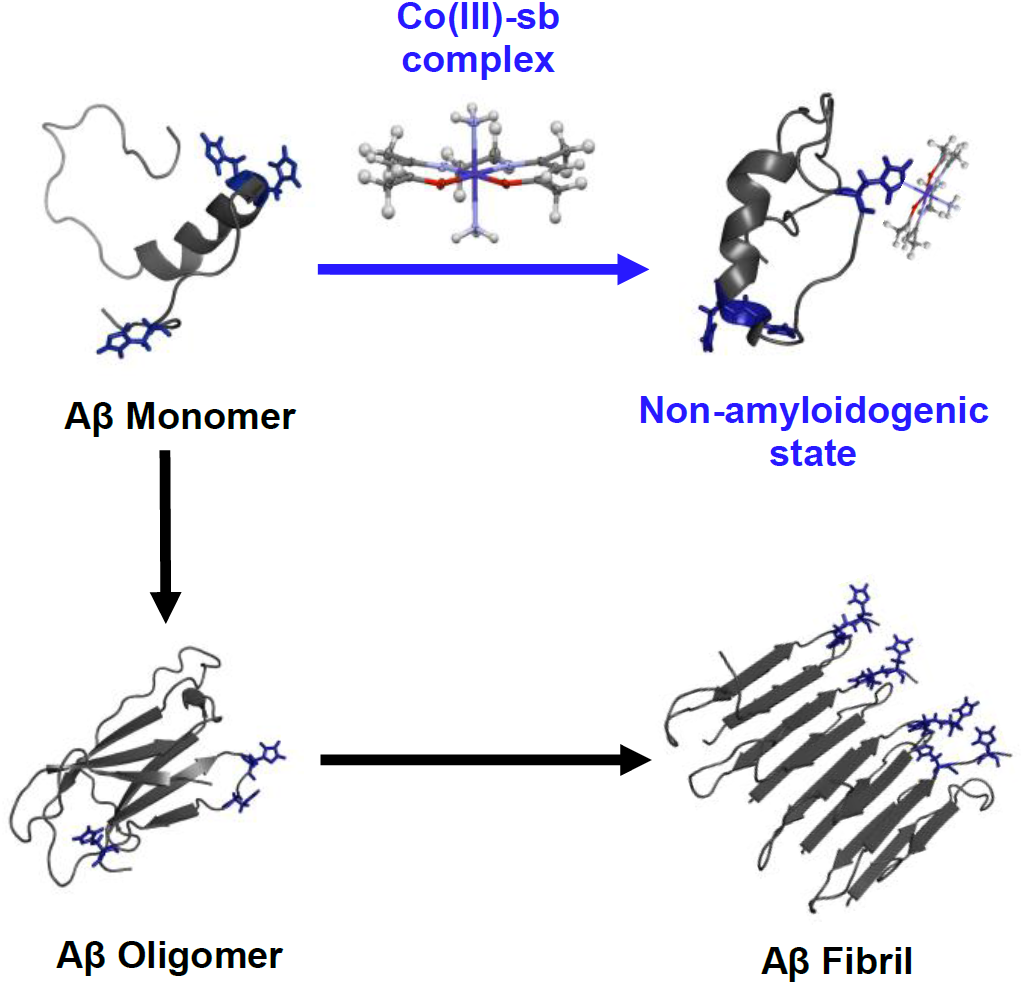
Co(III)-sb complexes bind monomeric Aβ through its N-terminal Histidine residues altering the aggregation kinetics and favoring the formation of non-amyloidogenic species. This results in the generation off-pathway Aβ oligomers that exhibit minimal β-sheet content and reduced cytotoxicity compared to untreated Aβ aggregates.

To this end, we have investigated the use of Co(III)-sb complexes toward the inhibition of Aβ aggregation.(27-29) Coordination of Co(III)-sb complexes altered the structure of Aβ_42_ peptides and promoted the formation of large soluble oligomers (**Scheme 1**). Interestingly, this structural perturbation of Aβ correlated to reduced synaptic binding to hippocampal neurons.(27) Further, computational studies using DSSP demonstrated the ability of Co(III)-sb to disrupt β-sheet formation in Aβ monomers and pentamers.(28, 29)

Here, we investigate how Co(III)-sb alters aggregation kinetics by a variety of complementary modalities including ThT fluorescence, CD spectroscopy, TEM, and AFM. We have performed kinetic fitting (AmyloFit) to gain insights into how Co(III)-sb alters Aβ aggregation and demonstrated that Co(III)-sb treated Aβ aggregates show reduced cytotoxicity to HT22 cells in culture.

## Materials and Methods

### Thioflavin T Fluorescence

As in Iscen et al (2019), Aβ_40_ and Aβ_42_ (AS-72216) were purchased from Anaspec (Fremont, CA).(28) This peptide was pretreated to ensure monomerization and came with a sample aggregation curve for an estimation of aggregation kinetics. The peptide was resuspended, sonicated for 30 seconds in a bath sonicator, and diluted in 20 mM HEPES, 100 mM NaCl to a final concentration of 50 μM Aβ_42_. A fresh solution of ThT was prepared for each assay and syringe filtered before use. ThT was added to the peptide mixture at a final concentration of 200 μM. Co(III)-sb and unmetallated acacen ligand (negative control) were synthesized and purified according to literature protocols and were added to the samples at concentrations ranging from 5 to 25 μM (0.1, 0.3, and 0.5 equivalents).^27^ Negative controls consisting of ThT and Co(III)-sb were run to ensure that Co(III)-sb does not alter ThT fluorescence. Samples were plated in quadruplicate on 384 well black, clear flat bottom polystyrene non-binding surface plates (Corning 3544). All peptide manipulation was performed as quickly as possible to ensure that early time points during the lag phase were not missed since aggregation proceeded very quickly. Typical preparations from resuspension to plating took 10 minutes.

Plates were sealed with aluminum plate seals to prevent sample evaporation. All peptide manipulation was performed using non-binding tips to avoid protein loss. Fluorescence was measured through the plate bottom every 5 minutes for 2 hours at excitation 440 nm and emission 484 nm on a Synergy H1M plate reader. Samples were kept at 37 °C under orbital agitation (252 rpm) in the plate reader between reads. Kinetic data from two separate plates was pooled and plotted using Graphpad Prism. Error bars represent standard error of the mean (SEM). Area under the curve was calculated for each replicate and data is presented as mean ± SEM.

### Circular Dichroism Spectroscopy

For circular dichroism studies synthetic lyophilized Aβ_40_ and Aβ_42_ (No. RP10017) were purchased from Genscript (Piscataway, NJ). This peptide was not guaranteed monomeric, so underwent a monomerization protocol before use. The lyophilized peptide was resuspended in 1,1,1,3,3,3 hexafluoroisopropanol for monomerization. Peptide mixtures of Aβ_40_ and Aβ_42_ were made by combining the appropriate amounts of each monomeric isoform in HFIP so that all aliquots contained 0.1 mg/tube. In order to test the effect of peptide isoform, four types of aliquots were produced: 100% Aβ40, 90% Aβ40 10% Aβ42, 50% Aβ40 50% Aβ42, and 100% Aβ42. The monomeric peptide was aliquoted into non-binding surface Eppendorf tubes at 0.1 mg/tube and allowed to dry under nitrogen for 24 hours, followed by 2 hours under vacuum to ensure removal of residual solvent. Following formation of dry peptide films, the aliquots were stored in a -80 °C freezer until use. All solutions used for experiments were syringe filtered through a 0.2 μm filter to remove particulates that could trigger aggregation.

For resuspension, dried peptide films were brought to room temperature and 50 μL of 20 mM NaOH was added to 0.1 mg of the peptide film. This solution was then probe sonicated for 6 x 1 second pulses at 20% amplitude. Following sonication the peptide was diluted into PBS containing 200 μM EDTA for chelation of residual metal ions and 0.02% sodium azide to inhibit bacterial growth. Aggregation was measured using peptide at a final concentration of 68 μM with Co(III)-sb at 0.5 eq with respect to protein (34 μM). All peptide manipulation was performed using non-binding tips to avoid protein loss.

Samples were read in a 1 mm quartz cuvette on a Jasco J-815 CD spectrometer. Reads were accumulations of 3 scans between 200 nm and 400 nm. Data pitch was set to 1.0 nm, bandwidth was 1 nm, and scan speed was 10 nm/min. Between reads the peptide was kept on an incubator shaker at 37 °C and agitated at 100 rpm. Molar ellipticity at 218 nm was be used to monitor aggregation kinetics over time. Data was analyzed using Jasco SpectraAnalysis software. Processing was completed using CDProAnalysis software.

### Transmission Electron Microscopy

Aggregated Aβ was characterized morphologically using transmission electron microscopy (TEM). Aggregated solutions were diluted to 10 μM and 10 μl was spotted onto formvar/carbon coated nickel grids (Electron Microscopy Sciences, Hatfield, PA) for 5 minutes. Excess solution was wicked off using filter paper. Grids were washed with deionized water twice for 1 minute before being stained with 10 μl 1% uranyl acetate for 5 minutes. Grids were dried overnight and imaged on a Hitachi HD-2300 Dual EDS Cryo STEM operated at 120 kV. Structures formed in the presence and absence of Co(III)-sb were qualitatively assessed for differences. Images were processed using ImageJ.

### Atomic Force Microscopy

Samples for Atomic Force Microscopy (AFM) were prepared by diluting 10 μl of the aggregation mixtures in 1 ml of double distilled water. 15 μl of each sample was then drop cast onto freshly cleaved mica plates (Ted Pella, Redding, CA) functionalized with (3-aminopropyl)triethoxysilane) (APTES). The samples were incubated at ambient temperature for 10 minutes, washed once with 1 ml of double distilled water, then allowed to dry protected from dust for at least 6 hours. APTES functionalized mica plates were produced by adding 10 μl of a 0.1 (v/v) aqueous solution of APTES to freshly cleaved mica and incubating at ambient temperature for 2 minutes. The plates were rinsed with 3 x 1 ml of double distilled water and allowed to dry, protected from dust, for at least 12 hours.

The samples were imaged using a Bruker Dimension FastScan system equipped with a silicon tip (nominal radius 10 nm). Initial image flattening and preparation of publication quality images were conducted in the NanoScope Analysis software package (v1.9). Equivalent disc radius and maximum bounding dimensions were calculated using Gwyddion analysis software (v. 2.50). The distributions of these values were calculated using an image mask with a minimum z-value of 0.5 nm. Data were plotted in Graphpad Prism and represent mean ± SEM.

### Kinetic Fitting using AmyloFit

Data obtained from ThT aggregation experiments was uploaded to the free online platform AmyloFit for processing and analysis of protein aggregation kinetic data.^9(30)^ AmyloFit is a free platform that can be accessed at http://www.amylofit.ch.cam.ac.uk/login. Mathematical fitting to protein aggregation models allowed determination of which microscopic processes were dominant and how treatment with Co(III)-sb affected these processes. Experiments were carried out as described above, with the exception that multiple dilutions of final Aβ monomer concentration were used (40 μM, 20 μM, 10 μM, and 5 μM). These dilutions were made by serially diluting the starting peptide concentrations down in buffer containing 200 μM ThT. Co(III)-sb dosing was always maintained at the aforementioned equivalencies compared to peptide concentration. ThT concentration was 200 μM for all monomer concentrations.

Raw data was reformatted to text files containing time and raw fluorescence measurements for each condition. Data was uploaded, normalized to 0% and 100%, and replicates were grouped. Fitting was attempted for untreated, 0.1 eq Co(III)-sb, 0.3 eq Co(III)- sb, and 0.5 eq Co(III)-sb treated conditions. Only untreated Aβ and Aβ + 0.1 eq Co(III)-sb were able to be fit by the program. The half time plotter tool was used to confirm the most appropriate model to fit the data. Next, the data was fit to the Nucleation Elongation model. The variables P_0_ and M_0_ were set to a Global Constant of 0. The variable m_0_ was set to a Constant for each data set and was equal to the starting monomer concentration for each aggregation. The parameters k_n_, n_c_, k_+_, and k_off_ were set to Global Fit with initial guesses of 1, 2, 1 x 10^8^, and 1 × 10^−15^ respectively. Basin hops was set to 50 and the data was fit with errors to yield the model parameters described below.

### AmyloFit Modeling

AmyloFit has the capability for modeling a variety of different aggregation mechanisms. The simplest of these is the nucleation elongation model. This model assumes that primary nucleation (ie. association of monomers into oligomeric nuclei) is the dominant method for formation of aggregation nuclei. This is in contrast to other models emphasizing the role of secondary nucleation (wherein larger aggregates can act as nuclei for the formation of new fibrils). The appropriateness of the model is determined by plotting the log of the initial monomer concentration (m_0_) vs the log of the aggregation half time (τ). Prior normalization of the data is critical for this step.^29^

The half time log plots demonstrate a linear relationship between the log of the initial monomer concentration and the log of the half time (**SI Figure 1**). The slope of this graph is termed the scaling exponent and contains information about the dominant mechanism of fibril multiplication. In addition, the shape of the curve offers insight into how the dominant mechanism changes with monomer concentration. A linear graph indicates that the dominant mechanism is the same for all monomer concentrations. As positive curvature in the graph indicates competition between two processes in series or a saturation process. A negative curvature indicates competition between two processes in parallel (for example both secondary nucleation and fibril fragmentation).^29^

### MTS Toxicity

HT22 cells were grown in DMEM media. 500 ml base media was supplemented with 10% fetal bovine serum (FBS), 5 ml non-essential amino acids, 5 ml L-glutamine, and 5 ml sodium pyruvate. Cells were passaged at approximately 70% confluence using trypsin. Cellular viability was assessed using the CellTiter 96 AQueous One Solution Cell Proliferation Assay (Promega, Madison, WI). Cells were plated in 96-well tissue culture treated plates at a density of 5,000 cells/well. Once cells had adhered, agents were dosed in replicate into the media at the desired concentrations. Viability was generally measure at 24 hours unless specified otherwise. After 24 hours, the media containing the agent was aspirated and replaced with 100 μl fresh media. 20 μl of 50% CellTiter solution in PBS was added to each well. Cells were incubated with the MTS reagent until signal was in the linear range (0.1 to 1 absorbance). 75 μl of the media containing the MTS reagent was then transferred to a new plate (to avoid absorbance error caused by cells on the bottom of the plate). Absorbance was measured at 490 nm on a Synergy H1M plate reader.

## Results and Discussion

### Co(III)-sb Alters Aggregation Kinetics by ThT Fluorescence

ThT fluorescence remains the most widely used method for measuring *in vitro* amyloid aggregation kinetics. To explore the effects of treatment with Co(III)-sb, monomeric Aβ was treated with 0.1, 0.3, and 0.5 equivalents of Co(III)-sb. Because each monomer of Aβ has three N-terminal His residues, the highest number of bound Co(III)-sb possible would be 3 equivalents assuming one to one binding with each His residue. As such, all doses used in these aggregation studies were substoichiometric for Co(III)-sb compared to Aβ. Co(III)-sb treated Aβ was compared to untreated control aggregation, as well as Aβ treated with the highest dose (0.5 equivalents) of the unmetallated ligand acacen (control).

Treatment with Co(III)-sb showed dose-dependent reduction in the aggregation by ThT fluorescence (**Figure 1**). For Aβ_42_, this was reflected in a decrease in the equilibrium plateau height, as well as a decrease in the slope of the curve during the polymerization phase, which manifests as an increase in the half time of aggregation (time to reach half max fluorescence). Explicit effects on the lag phase (initial period of baseline signal) were obscured by the rapid kinetics of aggregation. Based on the sample aggregation curve sent with the protein samples, the lag phase of aggregation was between 5 and 10 minutes. Due to sample prep, the first read was not until 10 minutes after resuspension, by which time the conditions had already entered into the polymerization phase of aggregation.

**Figure 1:**
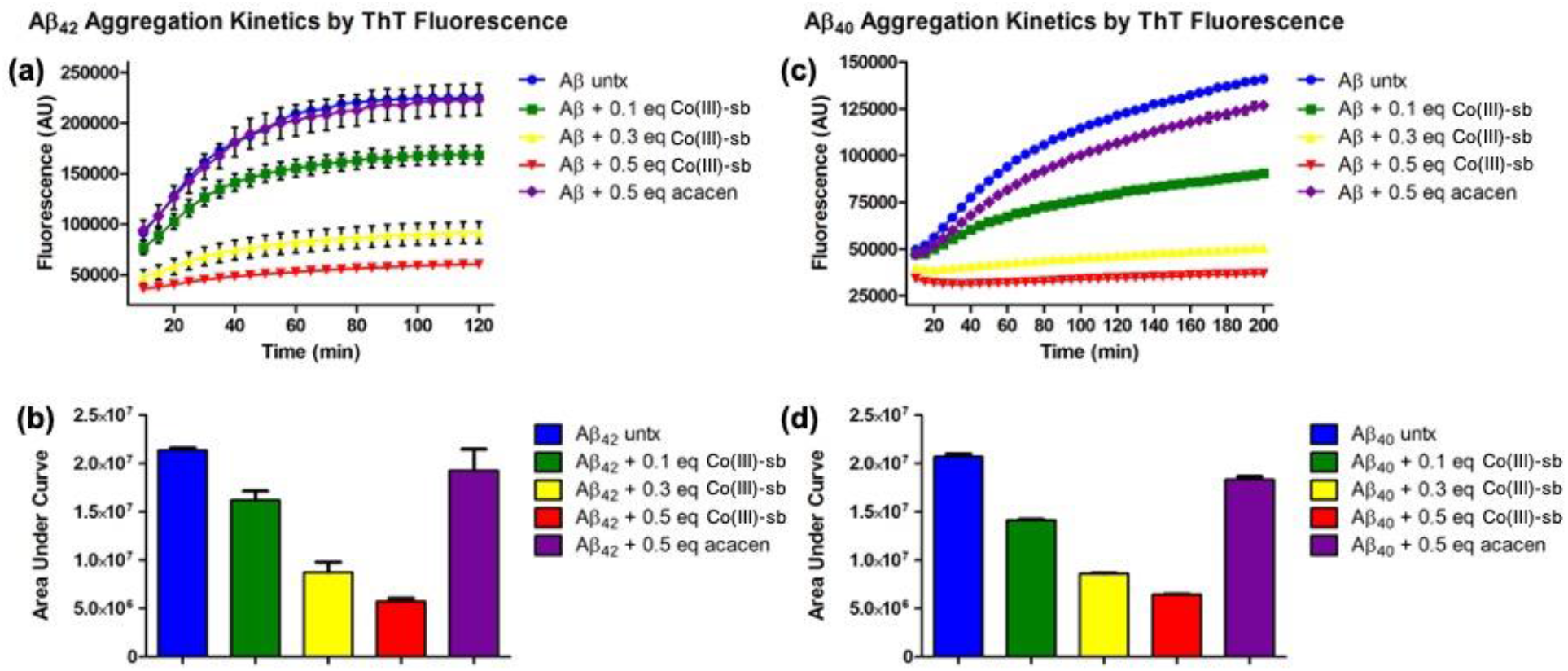
Inhibition of Aβ_42_ (a) and Aβ_40_ (c) aggregation by Co(III)-sb by ThT fluorescence. AUC analysis of Aβ_42_ (b) and Aβ_40_ (d) aggregation by ThT fluorescence. Untreated Aβ is in blue, Aβ + 0.1 eq Co(III)-sb is in green, Aβ + 0.3 eq Co(III)-sb is in yellow, Aβ + 0.5 eq Co(III)-sb is in red, and Aβ + 0.5 eq acacen is in purple. Data are presented as mean ± SEM. Co(III)-sb demonstrates dose-dependent inhibition, while acacen (negative control) does not alter aggregation.

Area under the curve analysis combines the effects on lag time and equilibrium height into a single metric (**Figure 1b**). This demonstrates that 0.1, 0.3, and 0.5 equivalents Co(III)-sb decreases the area under the aggregation curve by 25%, 60%, and 68% respectively compared to untreated control. In contrast, treatment with 0.5 equivalents of the unmetallated ligand (control) in does not significantly alter the area under the aggregation curve.

Repeating the same aggregation studies using Aβ_40_ instead of Aβ_42_ exhibited similar results (**Figure 1c**). Co(III)-sb exhibits dose-dependent inhibition of aggregation with a reduction in both equilibrium height and polymerization rate. Additionally, 0.5 equivalents of Co(III)-sb is enough to completely inhibit aggregation of the peptide. With Aβ_40_, the aggregation kinetics are slower that previously described for Aβ_42_, but are still on the faster end of kinetics reported in the literature. In addition, the negative control (0.5 equivalents acacen) appears to have a small inhibitory effect on the Aβ_40_ aggregation, which was not the case for Aβ_42_.

AUC analysis for Aβ_40_ is very similar to that for Aβ_42_, with 0.1, 0.3, and 0.5 equivalents of Co(III)-sb inhibiting the aggregation by 31%, 57%, and 67% respectively (**Figure 1d**). Treatment with 0.5 equivalents of acacen decreased aggregation by only 12%, indicating that although the negative control demonstrated some inhibition, it was to a much smaller degree than even one fifth the dose of the metallated active compound. This data is in agreement with our previously published aggregation studies using an alternate form of Co(III)-sb with methylamine axial ligands (instead of the bisammine complex utilized in this study). Direct comparison of the two complexes suggests that slightly enhanced aggregation inhibition is achieved with the bisammine Co(III)-sb used in this paper.(28)

### Co(III)-sb Mechanistic Insight from Kinetic Fitting

After significant optimization effort, a robust protocol for measuring Aβ aggregation by ThT fluorescence was produced that yielded high-quality reproducible kinetic curves. A major advantage for measuring full aggregation kinetics (in contrast to aggregation end point assays such as SDS-PAGE, AFM, or TEM) is that mathematical modeling can be used to extract mechanistic details about the mode of inhibition. Traditionally, aggregation curves have been fit using sigmoidal equations suited to mathematically describe the shape of the curve.(30) While useful for comparing the effects of treatments on the shape of the overall curve, these equations were not rooted in the underlying microscopic processes of aggregation. More recently, Knowles et al. have developed a system of derived rate laws describing the kinetics of amyloid aggregation that directly physically relates to the underlying protein association phenomena.(8) Knowles et al also developed an online platform called AmyloFit (available at http://www.amylofit.ch.cam.ac.uk/login), for facile fitting of outside generated aggregation data using their derived rate laws.(9)

In order to fit aggregation data using AmyloFit, aggregation kinetics must be generated at a variety of different initial monomer concentrations **(Figure S2)**. Linear plots of the aggregation half-life for each concentration are then constructed to determine aggregation mechanism (**Figure S1)**. The previously described ThT assay protocol was adapted to serially diluted protein monomer concentrations (40 μM, 20 μM, 10 μM, and 5 μM) while keeping ThT concentration constant. Co(III)-sb dosing was always expressed in molar equivalents compared to monomeric Aβ. Because the AmyloFit system was designed based on data from Aβ_42_, we used only kinetic data from Aβ_42_ aggregations. We attempted to model the results from all three tested doses of Co(III)-sb, however, 0.3 and 0.5 equivalents inhibited so strongly it rendered the curves non-sigmoidal which are incompatible with the mathematical model used by AmyloFit. As such,, and the results presented are limited to the comparison of the untreated Aβ_42_ to that treated with 0.1 equivalent of Co(III)-sb.

The results of the AmyloFit modeling technique can be found in **Table 1.**The most notable changes between the untreated Aβ_42_ and the Aβ_42_ + 0.1 equivalent Co(III)-sb were a 10 order of magnitude increase in the nucleation rate (k_n_) and a 9 order of magnitude decrease in the polymerization rate (k_+_) with Co(III)-sb treatment. All other parameters were either constants or did not vary significantly between the aggregation fits with and without Co(III)-sb. The variable k_off_ was very small for both data sets indicating that fibrils depolymerization played a very minor role in the aggregation kinetics and this was unchanged by Co(III)-sb.

**Table 1:** Global parameters from AmyloFit kinetic fitting of untreated Aβ_42_ aggregation and Aβ_42_ + 0.1 equivalent Co(III)-sb.

| Parameter | Untreated A$\beta_{42}$ | A$\beta_{42}$ + 0.1 eq Co(III)-sb |
| --- | --- | --- |
| P<sub>0</sub> (conc) | 0.00 | 0.00 |
| M<sub>0</sub> (conc) | 0.00 | 0.00 |
| $k_n$ ($\text{conc}^{-n_c+1} \text{ time}^{-1}$) | 5.72 E -8 (+1.7 E -8 / -1.5 E -9) | 230 (+16 / -22) |
| $n_c$ (unitless) | 0.614 (+1.1 E -2 / -4.0 E -3) | 0.693 (+2.5 E -4 / -7.8 E -3) |
| $k_+$ ($\text{conc}^{-1} \text{ time}^{-1}$) | 5.28 E 6 (+0.0 / -1.7 E 6) | 2.45 E -3 (+2.5 E -4 / -1.6 E -4) |
| $k_{\text{off}}$ ($\text{time}^{-1}$) | 6.94 E -15 (+3.1 E -16 / -1.4 E -15) | 2.29 E -15 (+0.0 / -9.3 E -16) |

### Co(III)-sb Aggregation Inhibition by CD Spectroscopy

CD spectroscopy is a complementary technique to ThT fluorescence that also follows aggregation kinetics by measuring β-sheet content in the protein as it aggregates. Aggregation mixtures were prepared as previously described for the ThT assay, but without the addition of exogenous fluorophore. Higher protein concentration was required for adequate signal, so dosing equivalency was maintained between assays. In order to obtain slow aggregation kinetics, Aβ_40_ was chosen as the isoform for these experiments.

Aβ_40_ was allowed to aggregate in a nonbinding Eppendorf tube and periodically transferred to a quartz 1 mm cuvette for measurement of ellipticity. When compared to the standard CD waveforms for various secondary structures, it is apparent that untreated Aβ_40_ initially existed in primarily a random coil structure, which was maintained throughout the first three timepoints (until hour 41). Between hours 41 and 68, the peptide underwent aggregation, shifting from a random coil waveform to a β-sheet waveform. Subsequent time points showed a stable β-sheet waveform that decreased in amplitude over time. This loss in signal was a result of protein precipitation to the cuvette, as well as loss of soluble protein aggregates.

As with the untreated Aβ_40_ aggregation, the 0.5 equivalent Co(III)-sb treated sample initially exhibited a random coil waveform at hour 1. However, while the random coil waveform was stable until hour 41 for the untreated, the Co(III)-sb treated Aβ_40_ quickly stabilized into an alternate conformation, not representing a random coil or β-sheet. This change was particularly interesting because it preceded the measured lag phase of aggregation for this Aβ_40_ prep (∼50 hours) and likely reflects a structural change during the nucleation phase of aggregation, which is very difficult to measure using ThT fluorescence.

Because Aβ aggregation represents a transition between two of the basic secondary structure waveforms (random coil to β-sheet), one way of quantifying this change to generate sigmoidal aggregation kinetics is to follow the ellipticity at 218 nm (the wavelength where the two curves differ the most). Aggregation is measured as a decrease in the signal at 218 nm indicating conversion to primarily β-sheet secondary structure. Like ThT aggregation, this method typically demonstrates a lag phase, a rapid polymerization phase, and an equilibrium plateau. **Figure 2c** illustrates sample kinetics for the previous data. Here it is obvious that the untreated Aβ_40_ undergoes a structural transition, while the Co(III)-sb treated Aβ_40_ does not.

**Figure 2:**
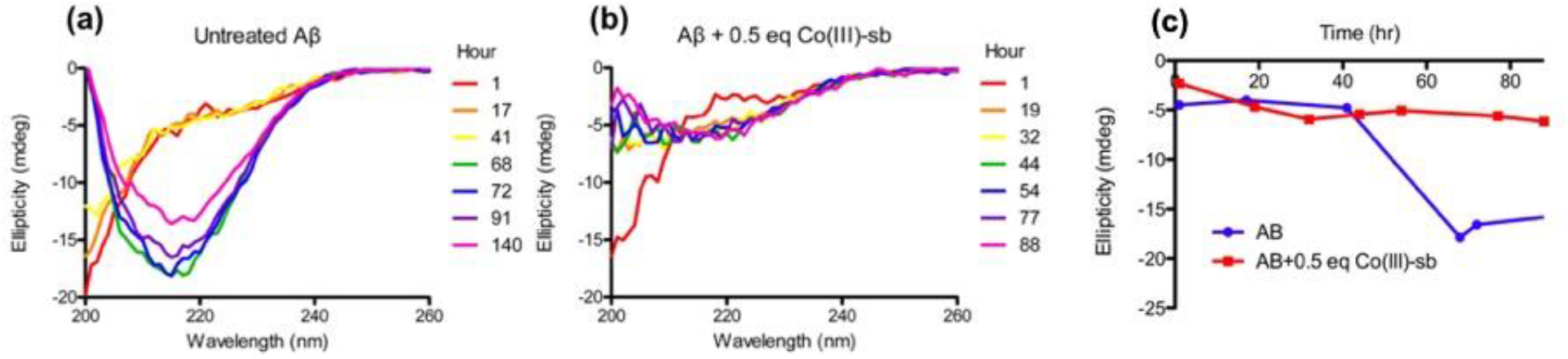
(a) Untreated Aβ_40_ aggregation by CD demonstrates a shift from random coil at early time points (red, orange, yellow) to β-sheet (green) after aggregation. (b) Aβ_40_ treated with 0.5 equivalents Co(III)-sb is initially random coil, but very early stabilizes into an alternative conformation. (c) Transition from random coil to β-sheet as measured by ellipticity at 218 nm. Untreated Aβ_40_ aggregates at around 50 hrs, while Co(III)-sb treated Aβ_40_ does not.

Additional CD experiments testing the other isoform ratios were conducted. Similar to ThT kinetics, it was observed that increasing the percentage of Aβ_42_ decreased the half time of aggregation. In addition, mixtures that had a high proportion of Aβ_42_ also precipitated out of solution at a higher rate. This led to low ellipticity signal, despite keeping protein concentrations constant. The signal for Aβ_42_ aggregation by CD was so low that the protein concentration required to see adequate signal was considered excessive. The trends described above for Aβ_40_ were additionally observed for the 9:1 mixture of Aβ_40_:_42_, but the absolute signal was lower (data not shown).

### Co(III)-sb Alters Aggregate Morphology by TEM

Structural changes in aggregate morphology were assessed via TEM. Grids spotted with the Aβ aggregation mixtures were negatively stained with uranyl acetate and imaged. **Figure 3a** demonstrates the formation of standard amyloid fibrils by untreated Aβ_40_. These fibrils are approximately 10 nm in diameter and up to μm in length. The fibrils form a dense meshwork and are negatively stained light on a dark background. In contrast, **Figure 3b** demonstrates the aggregates present following treatment with 0.5 equivalents Co(III)-sb. The aggregates are smaller than full fibrils and likely represent protofibrils or high molecular weight oligomers. Although not included in this figure, occasional fibrils were observed.

**Figure 3:**
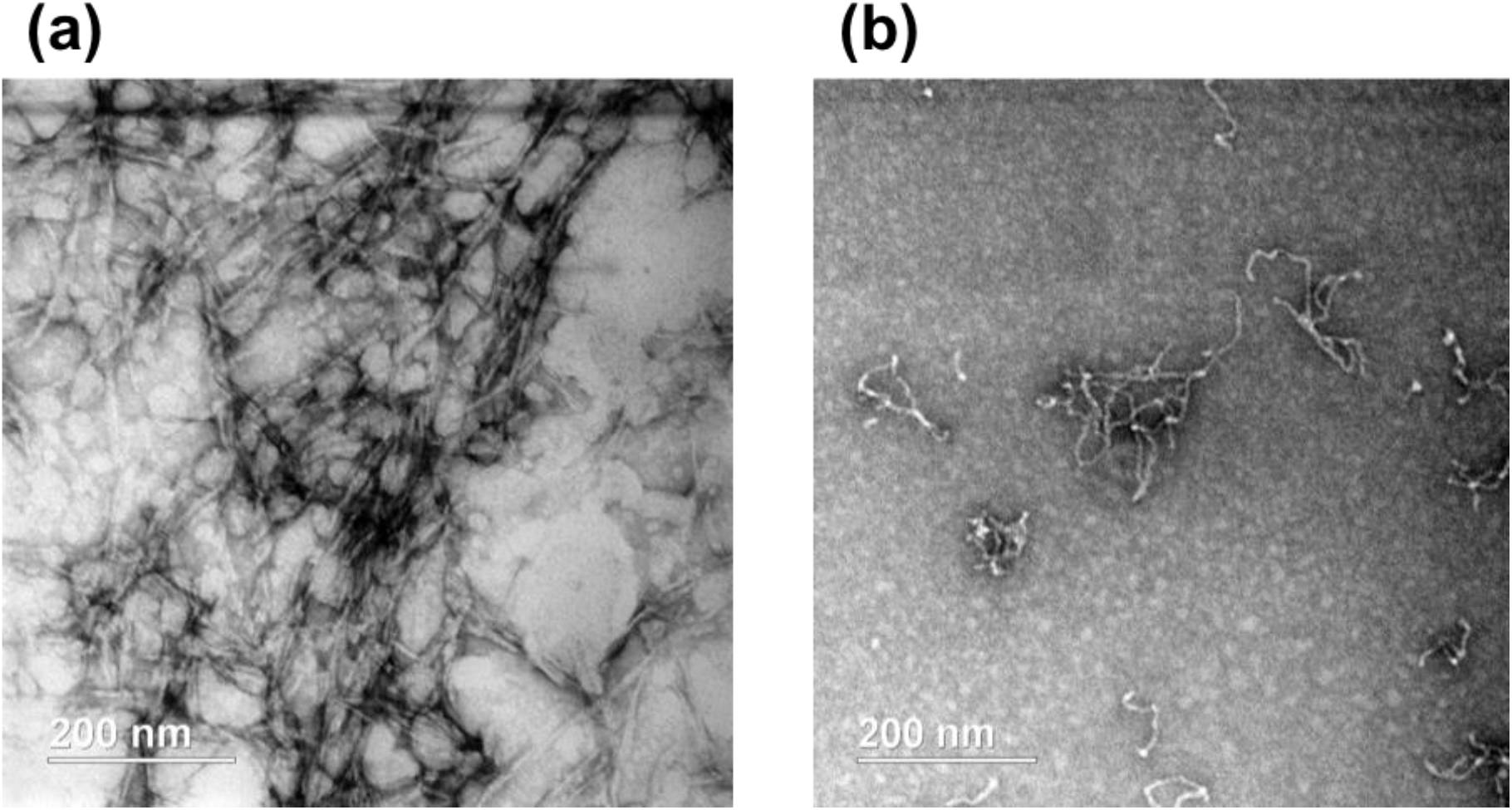
TEM images of (a) untreated Aβ_40_ demonstrating typical amyloid fibrils and (b) Aβ_40_ treated with 0.5 equivalents Co(III)-sb showing stabilization of smaller aggregate species morphologically consistent with protofibrils or high molecular weight oligomers.

TEM images were taken of Aβ_42_ aggregates with and without Co(III)-sb. In a similar fashion, the untreated protein demonstrated formation of typical amyloid fibrils (**Figure 4a**) which were shorter in length than those observed for Aβ_40._ This variation is likely due to underlying differences between the isoforms. The Co(III)-sb treated peptide showed a lack of fibrils (**Figure 4b**). Several small aggregate clusters were identified, but none of the spiral shaped winding protofibrils seen in the Aβ_40_ aggregations were evident in the Aβ_42_ samples.

**Figure 4:**
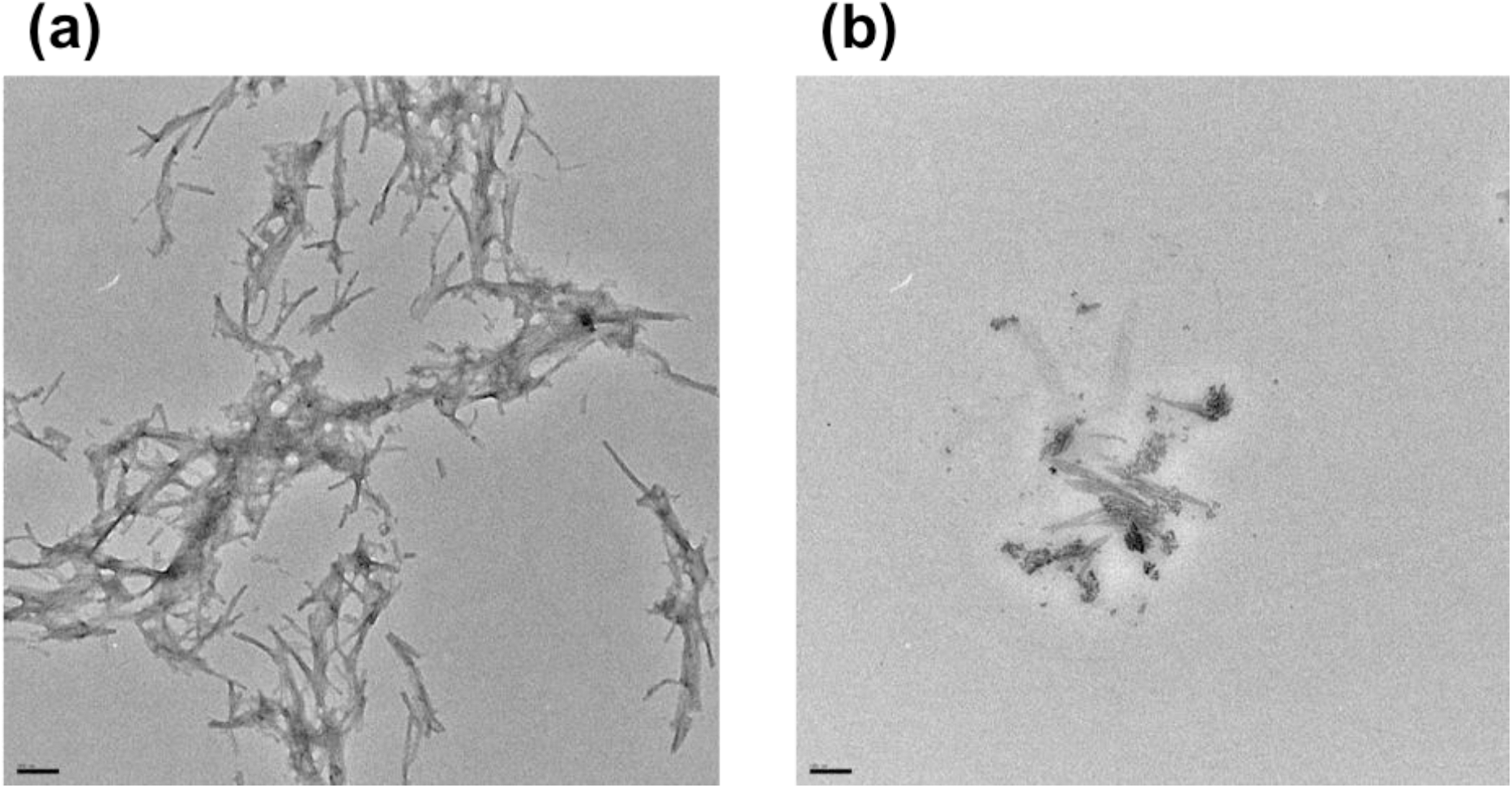
TEM images of (a) untreated Aβ_42_ showing typical amyloid fibrils and (b) Aβ_42_ treated with 0.5 equivalents Co(III)-sb showing stabilization of smaller aggregate species.

### Co(III)-sb Alters Size Distribution by AFM

AFM is a commonly used technique for morphological characterization of amyloid aggregates. It has higher resolution than TEM and is thus better suited to capturing effects on smaller aggregates and oligomers.(31) Equilibrium aggregation mixtures were diluted, spotted onto mica grids, and imaged. Comparison of particle sizes between untreated Aβ_42_ (**Figure 5a**), Aβ_42_ + 0.5 equivalents Co(III)-sb (**Figure 5b**), and Aβ_42_ + 0.5 equivalents acacen (**Figure 5c**) show that Co(III)-sb reduces the aggregate size, while acacen is not different from untreated.

**Figure 5:**
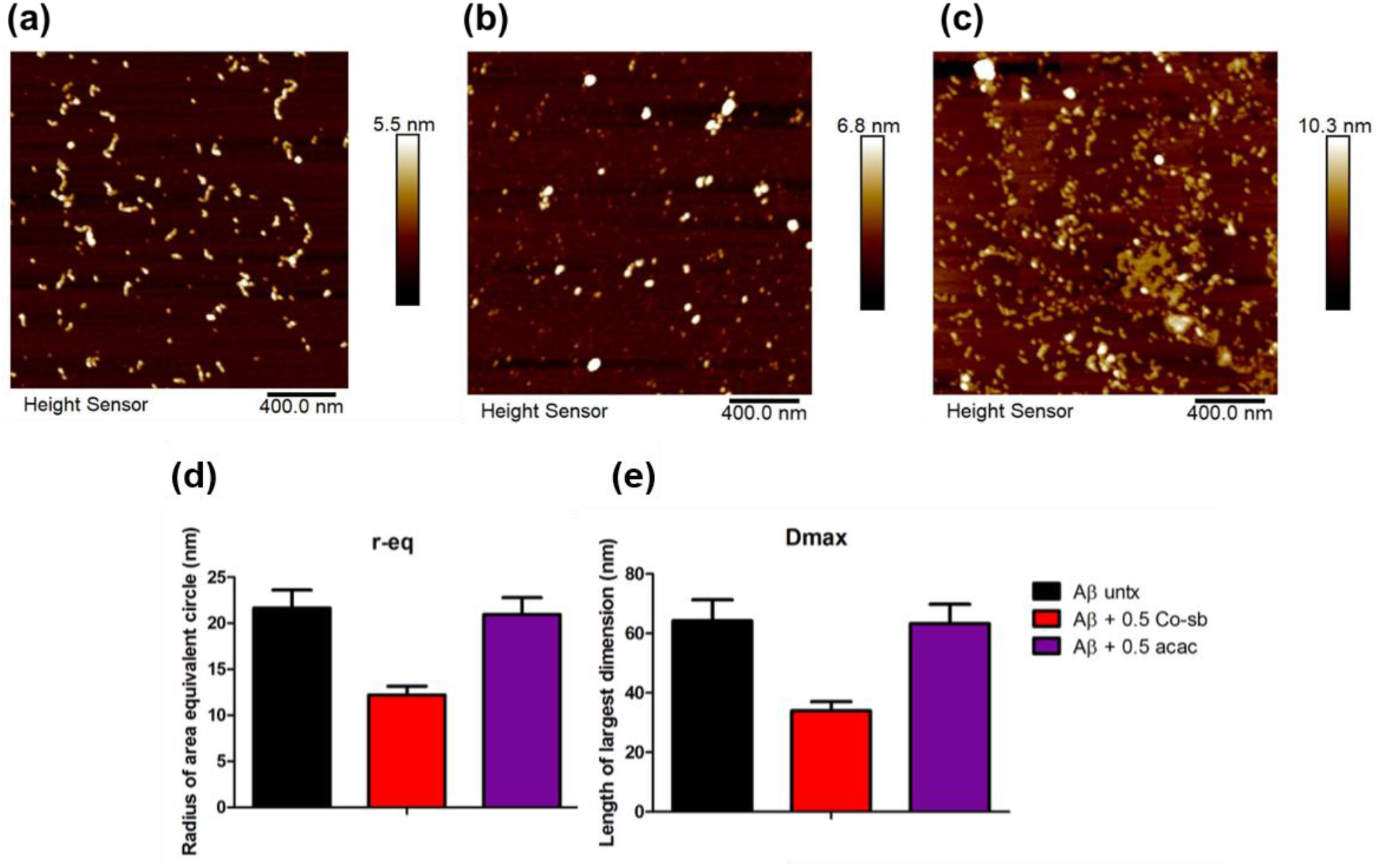
AFM images of equilibrium aggregate distributions for (a) untreated Aβ_42_, (b) Aβ_42_ + 0.5 equivalents Co(III)-sb, and (c) Aβ_42_ + 0.5 equivalents acacen (control). Quantitation of AFM size distributions. (d) Treatment of Aβ_42_ with Co(III)-sb decreases r-eq (the radius of the circle with equivalent area to aggregate size), while acacen does not differ from untreated Aβ_42_. (e) Treatment of Aβ_42_ with Co(III)-sb decreases Dmax (the largest dimension of aggregate size), while acacen does not differ from untreated Aβ_42_.

Quantitation of these particle size distributions is demonstrated in **Figure 5d and 5e**. The variable R-eq represents the radius of a circle with the same average area as the particles, while the variable D-max gives the average length of the largest dimension of the aggregates. In both area and length metrics, treatment with Co(III)-sb reduces the size of the aggregates, while treatment with the negative control acacen does not change the size. Additionally, the difference in D-max value is consistent with the difference longer, thinner amyloid fibrils compared to globular oligomers and protofibrils formed in the presence of Co(III)-sb.

### Toxicity Studies

Because many things are known to affect Aβ aggregation, it is especially important to demonstrate a biological endpoint when developing an amyloid inhibitor. As such we utilized cell viability assays to explore the effects of Co(III)-sb on Aβ-mediated toxicity to cultured cells. Co(III)-sb itself is not toxic to cells at the concentrations used for aggregation inhibition (**Figure S3**).

We then demonstrated rescue of cellular Aβ toxicity with Co(III)-sb treatment (**Figure 6**). 0.5 equivalents of Co(III)-sb, CoCl_2_, acacen alone did not alter viability **(Figure S3)**. Aggregated Aβ decreased cell viability to 70% of controls after 24 hours. Aβ aggregated in the presence of increased equivalents of Co(III)-sb showed increasing viability in a dose-dependent manner, while Aβ aggregated in the presence of the negative control acacen did not differ from untreated Aβ.

**Figure 6:**
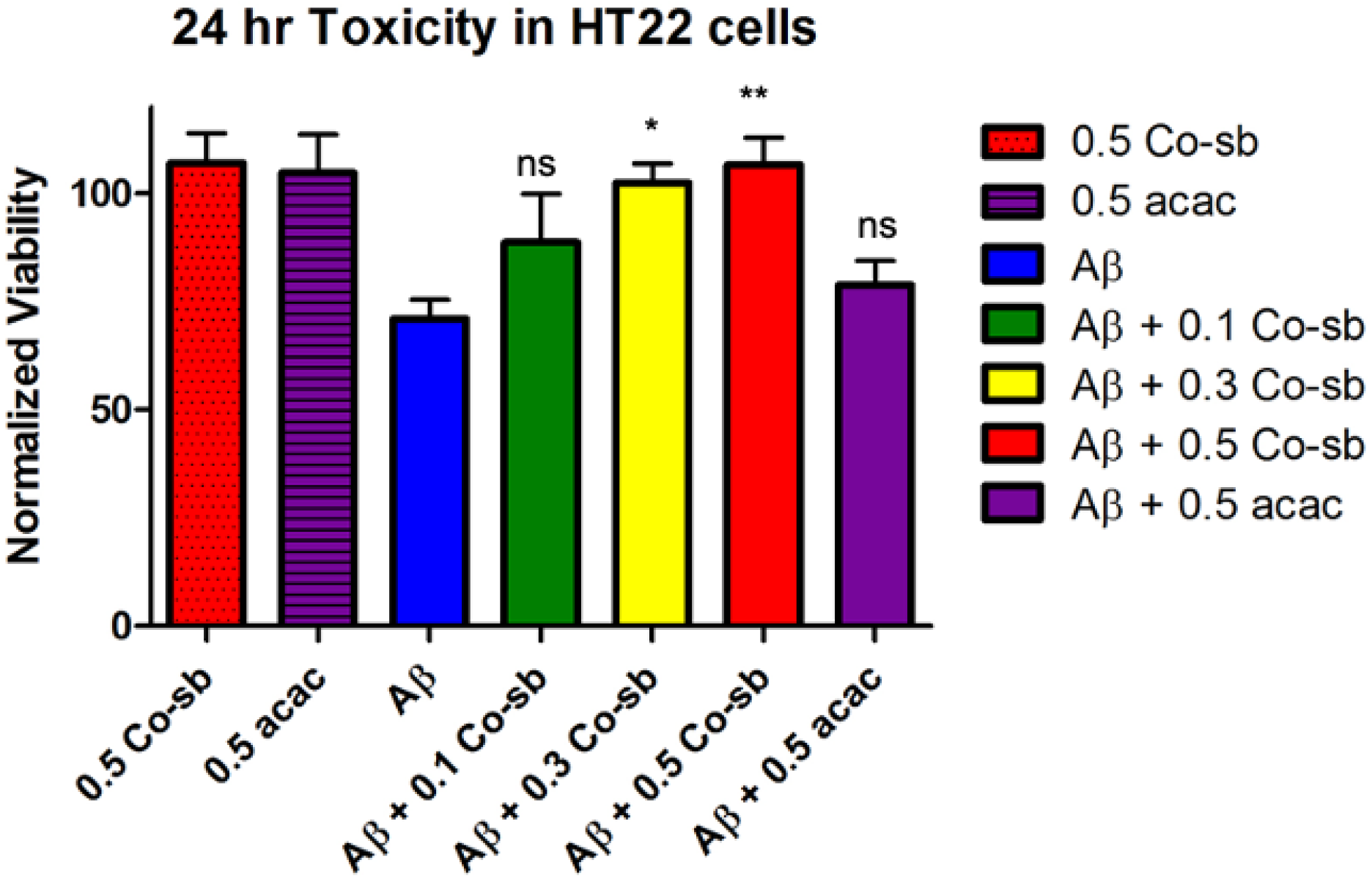
Rescue of Aβ toxicity by Co(III)-sb in HT22 cells by MTS assay. Treatment with Co(III)-sb reduced Aβ toxicity in a dose-dependent manner, while treatment with acacen had no effect. Data are normalized to the viability of vehicle treated controls and presented as mean ± SEM. * p < 0.05, ** p < 0.01.

ThT fluorescence assays showed robust dose-dependent inhibition of Aβ_40_ and Aβ_42_ aggregation. Further, this effect was specific to coordination by Co(III)-sb since the unmetallated negative control acacen did not inhibit aggregation. This inhibition was observed over a relatively narrow dosing window with 0.1 equivalent demonstrating moderate inhibition (∼30%) and 0.5 equivalents nearly completely inhibiting aggregation. Taken together, these data demonstrate potent inhibition of Aβ aggregation by Co(III)-sb.

## Conclusions

We have previously demonstrated through both computational and experimental studies that Co(III)-sb complexes can inhibit β-sheet formation during Aβ aggregation.(27-29) In this work we have expanded on these results by examining how Co(III)-sb binding alters the kinetic pathway of Aβ aggregation as well as characterized the secondary structure, morphology, and cytotoxicity of Co(III)-sb bound Aβ aggregates. Our kinetic assays employed ThT fluorescence which is a β-sheet sensitive fluorophore commonly used to study amyloid aggregation in a kinetic fashion. Despite its widespread use, there have been conflicting reports of ThT alone altering Aβ aggregation kinetics. Therefore, it is important to complement the results of ThT aggregation studies with complimentary techniques such as TEM, AFM, and CD which can detect the aggregation of Aβ without the use of an exogenous fluorophores. In addition to confirming the results of our ThT assays, AFM and TEM imaging demonstrated that Co(III)-sb complexes induce the formation of smaller, non-fibrillar oligomeric Aβ species. In particular, our CD results identified that changes to Aβ secondary structure caused by Co(III)-sb coordination happen early during the lag phase of the aggregation process, which could not have been observed by ThT fluorescence alone. This implies Co(III)-sb binding to Aβ likely alters the oligomeric distribution early in Aβ aggregation before the formation of β-sheet rich aggregates which are the species readily detected by ThT fluorescence.

The ThT kinetic data was fit to mathematical models of Aβ aggregation to determine the underlying mechanism of Co(III)-sb inhibition. According to the model, treatment with Co(III)-sb drastically increased the nucleation rate, and substantially decreased the polymerization rate. This corresponds to Co(III)-sb coordination to Aβ occurring early in the aggregation process and resulting in the rapid formation of oligomers that do not polymerize into larger aggregates or mature fibrils.

Cytotoxicity of the Co(III)-sb bound aggregates were tested using an immortalized rat hippocampal cell line (HT22) and showed reduced toxicity compared to untreated Aβ aggregates. Coupled with our previous work demonstrating Co(III)-sb treated aggregates exhibit reduced synaptic binding to primary hippocampal neurons this result suggests that Co(III)-sb drives the formation of oligomeric species which have weaker interactions with neuronal cell membranes, and therefore reduced cytotoxicity.(27)

Overall, Co(III)-sb complexes show great promise as anti-amyloid agents affecting both aggregation and toxicity. Direct comparison to other coordination complexes is difficult due to variation in assays, but a more detailed review of the many metal-based amyloid inhibitors can be found in Bajema et al. (2019) and demonstrates the broad utility of this class of complexes as amyloid inhibitors.(32) Future work will seek to further examine the molecular mechanisms of how Co(III)-sb alters aggregation, as well as further development of the complexes for use *in vivo*.

## Author Contributions

KFR and TJM designed the research. KFR, CRB, AP, DB, EH, and EV performed the experiments and data analysis. KFR, CRB, and TJM wrote and prepared the manuscript.

## Acknowledgments

This work was supported in part by the National Institutes of Health grant number R01GM121518. We thank Sara Fernandez from Northwestern University’s High Throughput Analysis Laboratory for assistance with ThT fluorescence assays, Eric Roth from Northwestern’s Electron Probe Instrumentation Center (EPIC) for help with TEM imaging, and Myrick Davis and Ron Soriano from Northwestern University’s Keck Biophysics Facility for their assistance with CD experiments.

This work was supported by the Northwestern University Keck Biophysics Facility and a Cancer Center Support Grant (NCI CA060553). This work made use of the EPIC facility of Northwestern University’s NU*ANCE* Center, which has received support from the Soft and Hybrid Nanotechnology Experimental (SHyNE) Resource (NSF ECCS-1542205); the MRSEC program (NSF DMR-1720139) at the Materials Research Center; the International Institute for Nanotechnology (IIN); the Keck Foundation; and the State of Illinois, through the IIN. This work made use of the SPID facility of Northwestern University’s NUANCE Center, which has received support from the Soft and Hybrid Nanotechnology Experimental (SHyNE) Resource (NSF ECCS-1542205); the MRSEC program (NSF DMR-1720139) at the Materials Research Center; the International Institute for Nanotechnology (IIN); the Keck Foundation; and the State of Illinois, through the IIN.

## References

1. 2020. 2019 - Alzheimer’s & Dementia. 2019 Alzheimer’s disease facts and figures Wiley Online Library, 2019 - Alzheimer’s & Dementia.

2. Selkoe, D. J., and J. Hardy. 2016. The amyloid hypothesis of Alzheimer’s disease at 25 years. EMBO Molecular Medicine 8(6):595–608.

3. Zempel, H., E. Thies, E. Mandelkow, and E. M. Mandelkow. 2010. Abeta oligomers cause localized Ca(2+) elevation, missorting of endogenous Tau into dendrites, Tau phosphorylation, and destruction of microtubules and spines. The Journal of neuroscience : the official journal of the Society for Neuroscience 30(36):11938–11950.

4. Viola, K. L., and W. L. Klein. 2015. Amyloid β oligomers in Alzheimer’s disease pathogenesis, treatment, and diagnosis. Springer Verlag. 183–206.

5. Forny-Germano, L., N. M. Lyra E Silva, A. F. Batista, J. Brito-Moreira, M. Gralle, S. E. Boehnke, B. C. Coe, A. Lablans, S. A. Marques, A. M. B. Martinez, W. L. Klein, J. C. Houzel, S. T. Ferreira, D. P. Munoz, and F. G. De Felice. 2014. Alzheimer’s disease-like pathology induced by amyloid-β oligomers in nonhuman primates. Journal of Neuroscience 34(41):13629–13643.

6. Nortley, R., N. Korte, P. Izquierdo, C. Hirunpattarasilp, A. Mishra, Z. Jaunmuktane, V. Kyrargyri, T. Pfeiffer, L. Khennouf, C. Madry, H. Gong, A. Richard-Loendt, W. Huang, T. Saito, T. C. Saido, S. Brandner, H. Sethi, and D. Attwell. 2019. Amyloid β oligomers constrict human capillaries in Alzheimer’s disease via signaling to pericytes. Science 365(6450):eaav9518.

7. Selkoe, D. J. 2001. Alzheimer’s disease: genes, proteins, and therapy. Physiol Rev 81(2):741–766.

8. Knowles, T. P., C. A. Waudby, G. L. Devlin, S. I. Cohen, A. Aguzzi, M. Vendruscolo, E. M. Terentjev, M. E. Welland, and C. M. Dobson. 2009. An analytical solution to the kinetics of breakable filament assembly. Science 326(5959):1533–1537.

9. Meisl, G., J. B. Kirkegaard, P. Arosio, T. C. Michaels, M. Vendruscolo, C. M. Dobson, S. Linse, and T. P. Knowles. 2016. Molecular mechanisms of protein aggregation from global fitting of kinetic models. Nat Protoc 11(2):252–272.

10. Cohen, S. I., M. Vendruscolo, C. M. Dobson, and T. P. Knowles. 2011. Nucleated polymerization with secondary pathways. II. Determination of self-consistent solutions to growth processes described by non-linear master equations. J Chem Phys 135(6):065106.

11. Cohen, S. I., M. Vendruscolo, C. M. Dobson, and T. P. Knowles. 2011. Nucleated polymerization with secondary pathways. III. Equilibrium behavior and oligomer populations. J Chem Phys 135(6):065107.

12. Cohen, S. I., M. Vendruscolo, M. E. Welland, C. M. Dobson, E. M. Terentjev, and T. P. Knowles. 2011. Nucleated polymerization with secondary pathways. I. Time evolution of the principal moments. J Chem Phys 135(6):065105.

13. Bartolini, M., M. Naldi, J. Fiori, F. Valle, F. Biscarini, D. V. Nicolau, and V. Andrisano. 2011. Kinetic characterization of amyloid-beta 1-42 aggregation with a multimethodological approach. Anal Biochem 414(2):215–225.

14. Bruggink, K. A., M. Muller, H. B. Kuiperij, and M. M. Verbeek. 2012. Methods for analysis of amyloid-beta aggregates. Journal of Alzheimer’s disease : JAD 28(4):735–758.

15. Grigolato, F., and P. Arosio. 2019. Sensitivity analysis of the variability of amyloid aggregation profiles. Physical Chemistry Chemical Physics 21(3):1435–1442.

16. Lee, M. C., W. C. Yu, Y. H. Shih, C. Y. Chen, Z. H. Guo, S. J. Huang, J. C. C. Chan, and Y. R. Chen. 2018. Zinc ion rapidly induces toxic, off-pathway amyloid-β oligomers distinct from amyloid-β derived diffusible ligands in Alzheimer’s disease. Scientific Reports 8(1):4772–4772.

17. Miura, T., K. Suzuki, N. Kohata, and H. Takeuchi. 2000. Metal binding modes of Alzheimer’s amyloid β-peptide in insoluble aggregates and soluble complexes. Biochemistry 39(23):7024–7031.

18. Collin, F., I. Sasaki, H. Eury, P. Faller, and C. Hureau. 2013. Pt(II) compounds interplay with Cu(II) and Zn(II) coordination to the amyloid-beta peptide has metal specific consequences on deleterious processes associated to Alzheimer’s disease. Chemical communications 49(21):2130–2132.

19. Faller, P., C. Hureau, and O. Berthoumieu. 2013. Role of metal ions in the self-assembly of the Alzheimer’s amyloid-beta peptide. Inorganic chemistry 52(21):12193–12206.

20. Faller, P., and C. Hureau. 2009. Bioinorganic chemistry of copper and zinc ions coordinated to amyloid-beta peptide. Dalton transactions(7):1080–1094.

21. Nam, E., G. Nam, and M. H. Lim. 2020. Synaptic Copper, Amyloid-β, and Neurotransmitters in Alzheimer’s Disease. Biochemistry 59(1):15–17.

22. Pithadia, A. S., and M. H. Lim. 2012. Metal-associated amyloid-β species in Alzheimer’s disease. 67–73.

23. Ryan, T. M., N. Kirby, H. D. T. Mertens, B. Roberts, K. J. Barnham, R. Cappai, C. L. L. Pham, C. L. Masters, and C. C. Curtain. 2015. Small angle X-ray scattering analysis of Cu <sup>2+</sup> -induced oligomers of the Alzheimer’s amyloid β peptide. Metallomics 7(3):536–543.

24. Mannini, B., J. Habchi, S. Chia, F. S. Ruggeri, M. Perni, T. P. J. Knowles, C. M. Dobson, and M. Vendruscolo. 2018. Stabilization and Characterization of Cytotoxic Aβ40 Oligomers Isolated from an Aggregation Reaction in the Presence of Zinc Ions. ACS Chemical Neuroscience 9(12):2959–2971.

25. Bush, A. I. 2013. The metal theory of Alzheimer’s disease. Journal of Alzheimer’s disease : JAD 33 Suppl 1:S277–281.

26. Barnham, K. J., V. B. Kenche, G. D. Ciccotosto, D. P. Smith, D. J. Tew, X. Liu, K. Perez, G. A. Cranston, T. J. Johanssen, I. Volitakis, A. I. Bush, C. L. Masters, A. R. White, J. P. Smith, R. A. Cherny, and R. Cappai. 2008. Platinum-based inhibitors of amyloid-beta as therapeutic agents for Alzheimer’s disease. Proceedings of the National Academy of Sciences of the United States of America 105(19):6813–6818.

27. Heffern, M. C., P. T. Velasco, L. M. Matosziuk, J. L. Coomes, C. Karras, A. L. Eckermann, W. L. Klein, and T. J. Meade. 2014. Modulation of Amyloid-beta Aggregation by Histidine-coordinating Co(III) Schiff Base Complexes. Chembiochem.

28. Iscen, A., C. R. Brue, K. F. Roberts, J. Kim, G. C. Schatz, and T. J. Meade. 2019. Inhibition of Amyloid-β Aggregation by Cobalt(III) Schiff Base Complexes: A Computational and Experimental Approach. Journal of the American Chemical Society 141(42):16685–16695.

29. Heffern, M. C., V. Reichova, J. L. Coomes, A. S. Harney, E. A. Bajema, and T. J. Meade. 2015. Tuning cobalt(III) Schiff base complexes as activated protein inhibitors. Inorganic chemistry 54(18):9066–9074.

30. Crespo, R., F. A. Rocha, A. M. Damas, and P. M. Martins. 2012. A generic crystallization-like model that describes the kinetics of amyloid fibril formation. The Journal of biological chemistry 287(36):30585–30594.

31. Ruggeri, F. S., T. Šneideris, M. Vendruscolo, and T. P. J. Knowles. 2019. Atomic force microscopy for single molecule characterisation of protein aggregation. Archives of Biochemistry and Biophysics 664:134–148.

32. Bajema, E. A., K. F. Roberts, and T. J. Meade. 2019. Cobalt-Schiff Base Complexes: Preclinical Research and Potential Therapeutic Uses. Met Ions Life Sci 19.

